# SGI-4 in monophasic *Salmonella* Typhimurium ST34 is a novel ICE that enhances resistance to copper

**DOI:** 10.1101/518175

**Authors:** Priscilla Branchu, Oliver J. Charity, Matt Bawn, Gaetan Thilliez, Timothy J Dallman, Liljana Petrovska, Robert A. Kingsley

## Abstract

A multi drug resistant (MDR) *Salmonella enterica* 4,[5],12:i-of sequence type 34 (monophasic *S*. Typhimurium ST34) is a current pandemic clone associated with livestock and numerous outbreaks in the human population. A large genomic island, termed SGI-4, is present in the monophasic Typhimurium ST34 clade and absent from other *S*. Typhimurium strains. SGI-4 consists of 78 open reading frames including *sil* and *pco* genes previously implicated in resistance to copper and silver, and multiple genes predicted to be involved in mobilisation and transfer by conjugation. SGI-4 was excised from the chromosome, circularised and transferred to recipient strains of *S*. Typhimurium at a frequency influenced by stress and induced by mitomycin C and oxygen tension. The presence of SGI-4 was associated with increased resistance to copper, particularly but not exclusively under anaerobic conditions. The presence of *silCBA* genes, predicted to encode an RND family efflux pump that transports copper from the periplasm to the external milieu, was sufficient to impart the observed enhanced resistance to copper. The presence of *silCBA* genes resulted in the absence of copper-dependent induction of a multicopper oxidase system encoded by *pco* genes, also present on SGI-4, suggesting that the system effectively limits the copper availability in the periplasm. The presence of *silCBA* genes did not affect SodCI-dependent macrophage survival.

## INTRODUCTION

*Salmonella enterica* serovar Typhimurium (S. Typhimurium) accounts for approximately 25% of all human cases of non-typhoidal *Salmonella* (NTS) infection in Europe, and it is widespread in multiple zoonotic reservoirs (1, 2). The epidemiological record of human multidrug-resistant (MDR) *S*. Typhimurium infections in Europe is characterized by successive waves of dominant MDR variants that persist for 10-15 years (3, 4). *S*. Typhimurium DT104 emerged around 1990, becoming a globally pandemic clone that affects numerous domesticated and wild animal species (5). In 2007 a monophasic *S*. Typhimurium (S. 4,[5],12:i-) variant emerged in European pig populations and spread globally; monophasic *S*. Typhimurium is multi-locus sequence type 34 (ST34) (hereafter referred to as monophasic *S*. Typhimurium ST34), and predominantly phage type DT193 or DT120 (6–12). The mechanisms for succession in *S*. Typhimurium variants in Europe are not known, but selection by commonly-used antibiotics is unlikely since variants share similar AMR profiles (3): ACSSuT (ampicillin, chloramphenicol, streptomycin, sulphonamide, tetracycline) for *S*. Typhimurium DT104 and ASSuT for monophasic *S*. Typhimurium ST34.

Copper is both an essential nutrient, due to its role as a cofactor in multiple enzymes in all aerobic organisms, and highly toxic due to its ability to displace iron from Fe-S complexes in dehydratases (13). To reduce toxicity, bacteria control the amount of free copper in the cytoplasm and periplasm using transport systems that oxidise cuprous (Cu^1+^) to cupric (Cu^2^+) ions (14). *Escherichia coli*, a close relative of *Salmonella*, encodes multiple transport systems on its chromosome to maintain copper homeostasis. *Escherichia coli* exports copper from the cytoplasm into the periplasm via: a P_1B_-type ATPase, CopA; a multicomponent copper RND family efflux pump, CusCFBA; and the multi-copper oxidase CueO. This safeguards the periplasm either by oxidation of the Cu or by transporting Cu to the outside of the cell (15, 16). CueO and CopA, which are co-regulated by the cytosolic CueR, are the primary copper homeostasis systems active during aerobic growth (18). In contrast CusCFBA is important during anaerobic growth and is under the transcriptional control of the periplasmic CusRS two component regulator (17). In addition, some *E. coli* isolated from copper-rich environments have additional plasmid-encoded genes *pcoABCDRSE*, that encode a multicopper oxidase system which is active in the periplasm and regulated by a two-component system with a sensor kinase that extends into the periplasm (14, 18).

Copper homeostasis in most strains of *Salmonella* is fundamentally different from that in *E. coli* because, although *Salmonella* species encode CopA and CueO, they normally lack the CusCFBA RND family efflux pump. This is due to deletion of the genes from the chromosome (19–21). Therefore, *Salmonella* species do not have the mechanism to transport Cu out of the cell entirely, which is likely to have a significant impact on the distribution of Cu within the cell. However, plasmid-borne *silRSCFB* genes have been described in a *S*. Typhimurium strain associated with an outbreak in burns patients that had been treated topically with silver nitrate; these genes encode an RND family efflux pump that is closely related to the CusCBA system and confers resistance to silver and copper (22, 23). The presence of *sil* or *cus* genes in *Salmonella* is uncommon although these genes have been reported in multiple distinct monophasic *S*. Typhimurium clones in the past two decades (11, 24).

When the whole genome sequence of monophasic *S*. Typhimurium ST34 was compared with other *S*. Typhimurium variants an 80kb genomic island was identified and designated as *Salmonella* genomic island 4 (SGI-4); this genomic island was present in over 95% of monophasic *S*. Typhimurium ST34 strains from the UK and Italy, but absent from a diverse collection of *S*. Typhimurium including DT104 (25). Ancestral reconstruction analysis was consistent with acquisition of SGI-4 by horizontal transfer concomitant with clonal expansion of monophasic *S*. Typhimurium ST34, and rare sporadic loss of the genetic island (25). SGI-4 contained two clusters of genes predicted to be involved in resistance to copper and silver or arsenic metal ions (25).

Here we addressed the hypothesis that SGI-4 is a mobile genetic element that encodes multiple metal ion resistance determinants that alter the growth of *Salmonella* in concentrations of copper relevant to the host and farm environment. Furthermore, we hypothesise that the copper homeostasis and silver resistance genes alter copper homeostasis and impact *Salmonella* survival in macrophages.

## RESULTS

### SGI-4 is a self-mobilizable member of a novel family of integrative conjugative elements (ICE)

We investigated the presence of candidate sequences within the SGI-4 coding proteins capable of mobilization to recipient bacteria. The mechanisms for transfer we considered are common to integrative conjugative elements (ICE), also known as conjugative self-transmissible integrative (CONSTIN) elements (26), integrative mobilizable elements (IMEs) or cis-mobilizable elements (CIMEs) (27). A number of ORFs exhibited sequence similarity to genes encoding DNA processing enzymes that are predicted to be involved in excision, integration and transfer of DNA. Type IV secretion systems (T4SSs) involved in conjugative transfer of DNA were also present on SGI-4 (Figure 1, Supplementary Table 2). We were unable to identify a putative oriT sequence in SGI-4 using sequence alignment to a database of known oriT sequences (28). However, two pairs of inverted repeats, each with 80% identity to one another, were present in ORF19 and in an intergenic region between ORF64 and ORF65.

**Figure 1.**
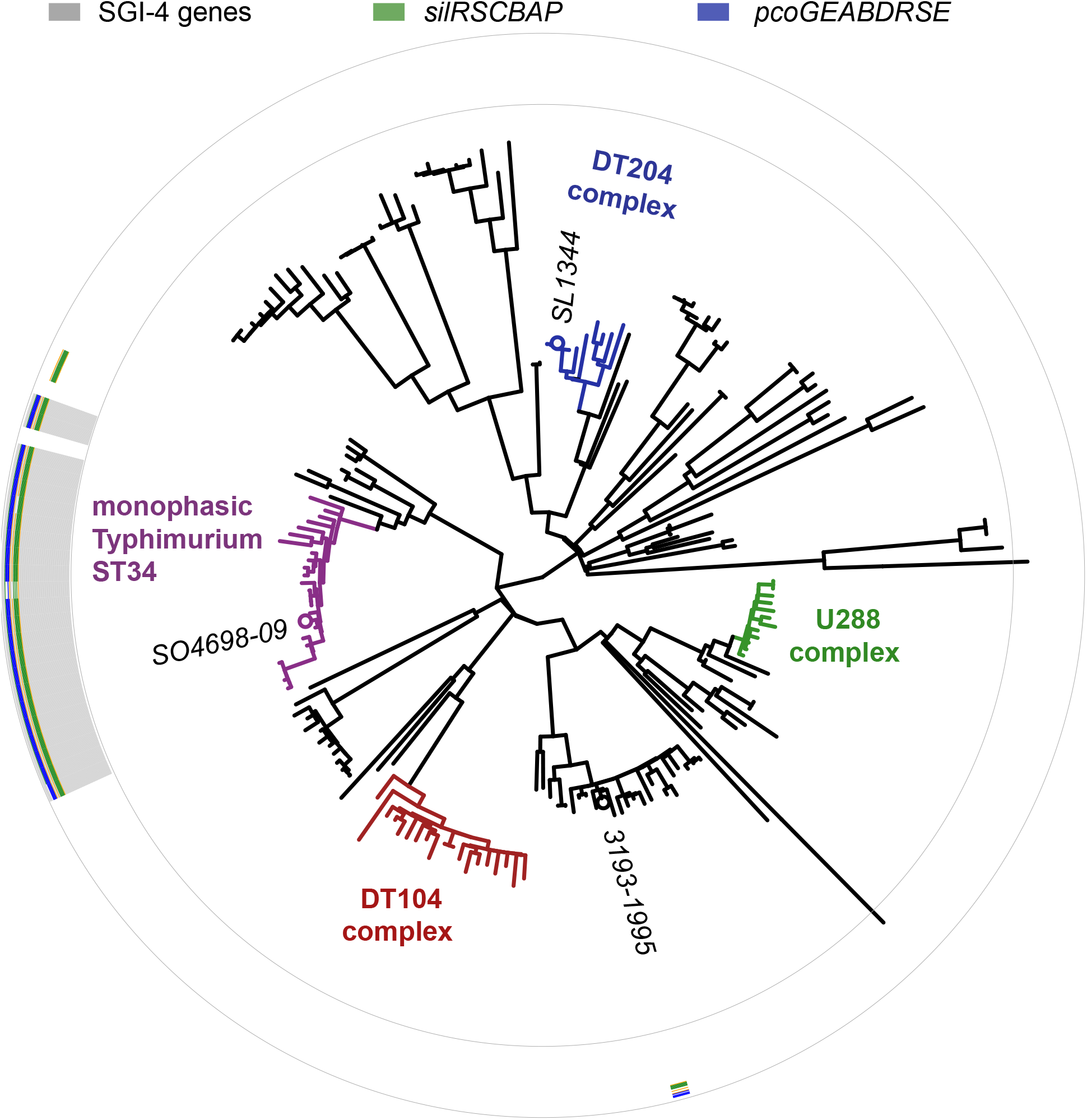
Distribution of SGI-4 in monophasic S. Typhimurium ST34 and representative strains of S. Typhimurium. A maximum likelihood tree constructed using sequence variation in the core genome of 24 monophasic *S*. Typhimurium ST34 strains and 153 representative strains of *S*. Typhimurium with reference to the whole genome sequence of strain SL1344 (accession FQ312003). The presence of sequences that mapped to 78 SGI-4 ORFs from SO4698-09 are represented as filled boxes in 78 concentric circles: *sil* genes (green), *pco* genes (blue), or other SGI-4 ORFs (grey).

**Figure 2.**
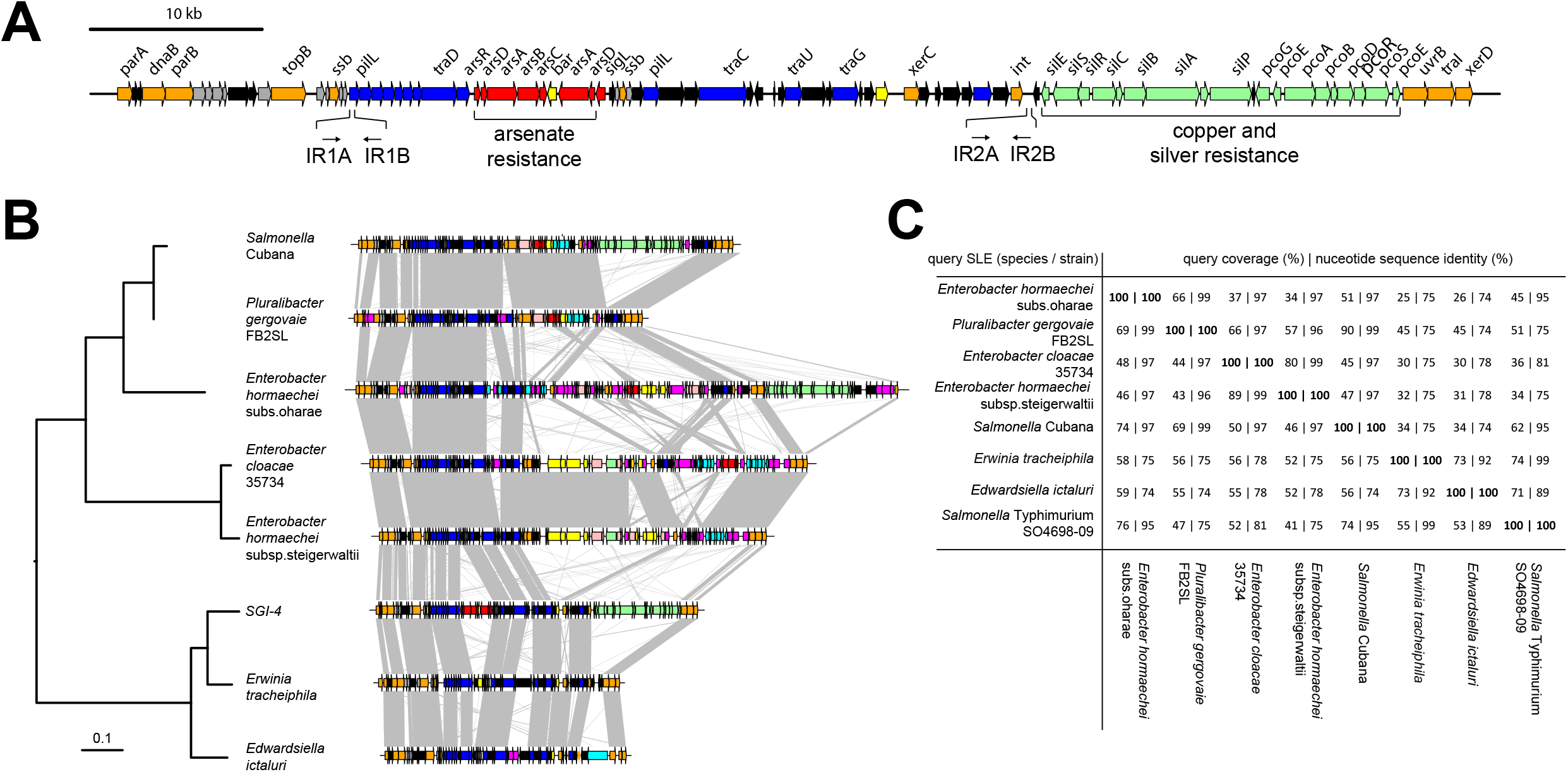
Genetic map of *Salmonella*. Genomic Island 4 (SGI-4) and phylogenetic relationship and gene synteny with SLEs. (A) Genetic diagram of *Salmonella* Genomic Island 4 (SGI-4); filled arrows indicate open reading frames with predicted functions based on sequence similarity for integration, excision and DNA processing (orange), type 4 secretion systems and conjugal transfer (blue), copper and silver resistance (light green), arsenic resistance (red), genes of unknown function commonly associated with ICE (grey), genes of unknown function not commonly associated with ICE (black). (B) Mid-point rooted maximum likelihood tree of eight SGI-4-like elements (SLEs) constructed using sequence variation in the core nucleotide sequence alignment of SLEs from diverse Enterobacteriaceae and genetic diagram of SLEs with regions of >80% nucleotide sequence identity (grey shading); filled arrows indicate open reading frames with the same designations as (A), with additional functions, cadmium, cobalt, zinc or mercury resistance (pink), transposase and insertion elements (purple), and a repressor protein lexA (brown).

To determine whether SGI-4 was similar to known MGEs we aligned the sequence of SGI-4 with nucleotide sequences in the NCBI database. Seven sequences in the NCBI database exhibited at least 75% nucleotide sequence identity with over 40% of SGI-4, none of which had had been described previously as MGEs (Figure 1B). We refer to these as SGI-4-like elements (SLEs). SLEs were inserted in the genome adjacent to a phenylalanine tRNA in strains from a diverse range of bacterial species from the Enterobacteriaceae, indicating this was the common attachment site (attB/attP). SGI-4 was most closely related to SLEs from *Erwinia tracheiphila* and *Edwardsiella ictaluri*, and more distantly related to SLEs from two *Enterobacter hormachei* strains from subspecies *oharae and steigerwaltii, Enterobacter cloacae, Pluralibacter gergoviae* and an SLE present in a strain of *Salmonella enterica* serotype Cubana (Figure 1B and 1C). All shared a number of regions of at least 75% sequence identity and synteny that encoded putative DNA processing and transfer functions (T4SS) and were interspersed amongst genes involved in other diverse functions including cargo genes capable of modifying the phenotype of the host bacterium. In several cases these genes were involved in resistance to a range of metals such as copper, silver, cadmium, mercury, zinc and cobalt (Figure 1B).

Since SGI-4 encoded many of the genes normally associated with ICE we determined whether it was capable of transferring a chloramphenicol resistance gene *(cat)* inserted into the arsenic resistance locus of SGI-4 to a recipient *S*. Typhimurium strain SL1344. Transfer frequency was low under aerobic culture conditions but was substantially increased in the presence of mitomycin C or by culture in anaerobic conditions (Figure 3). Mitomycin C and anaerobicity had an additive effect on transfer and, in combination, resulted in an increase in frequency of over four orders of magnitude. PCR amplification and sequence analysis of the junction site of the donor strain indicated that, on transfer, SGI-4 inserts into the same position on the chromosome in the *thrW* locus (data not shown). Furthermore, in the monophasic *S*. Typhimurium strain SO4698-09 cultured in the presence of mitomycin C, PCR amplification and sequence analysis of an amplicon generated using specific outward facing primers at each end of SGI-4, was consistent with circularisation of excised SGI-4 (data not shown).

**Figure 3.**
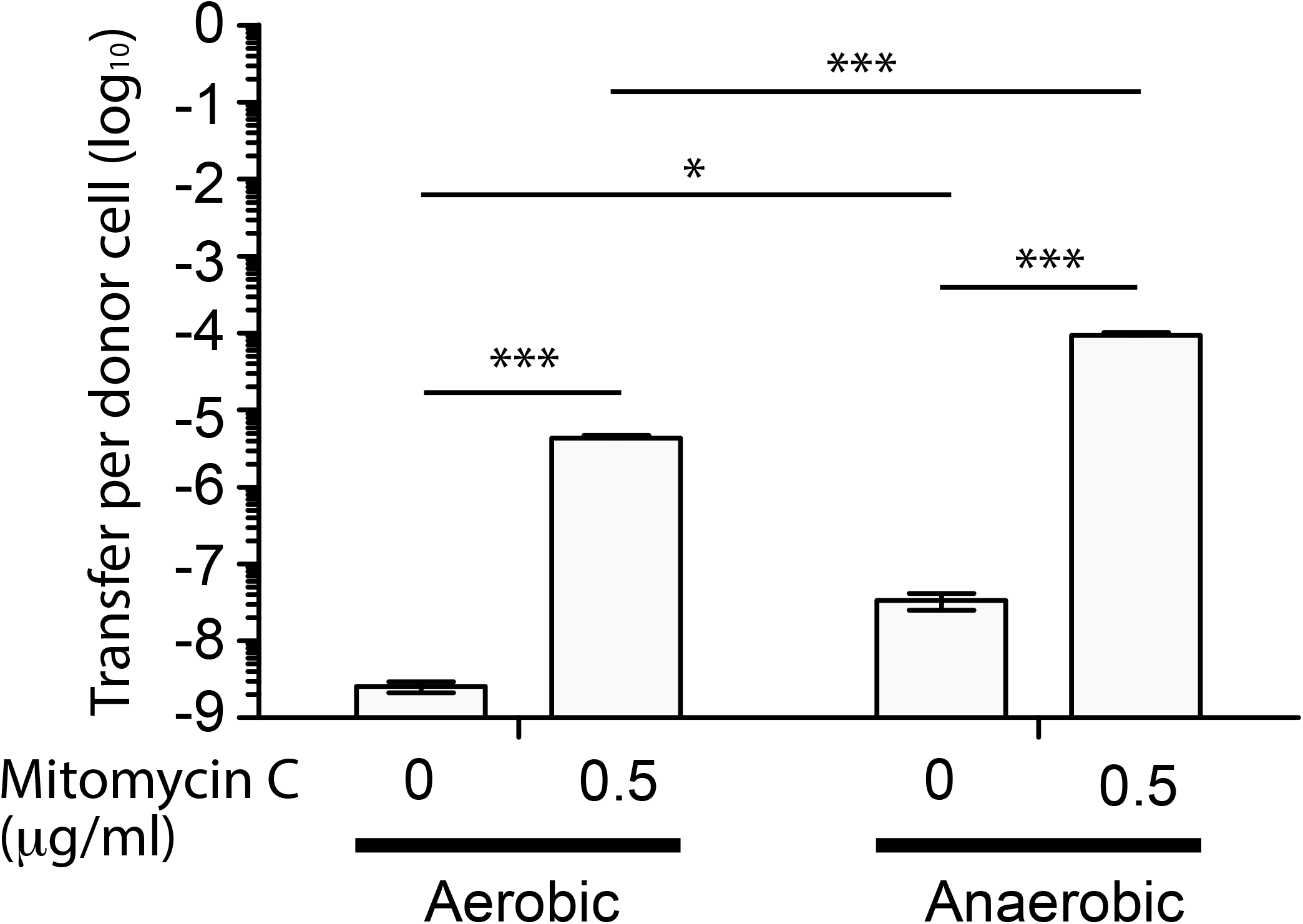
Transfer frequency of SGI-4::*cat* to S. Typhimurium during co-culture is enhanced by mitomycin C and hypoxia. Bars indicate the mean number of chloramphenicol resistant *S*. Typhimurium strain SL1344 colony forming units (CFU) per CFU of monophasic *S*. Typhimurium ST34 strain SO4698-09 SGI-4 *silCBA:: cat*, following co-culture. The mean from six biological replicates +/− standard deviation are shown.

### SGI-4 is characteristic of the monophasic *S*. Typhimurium ST34 clade

The SGI-4 sequence was present in 23 of 24 strains in the monophasic *S*. Typhimurium ST34 clade and absent from all other strains investigated, which were representative of common phage types of *S*. Typhimurium isolated from domesticated and wild animals and human clinical infections that occurred in the UK in the last 25 years (Figure 1 and Supplementary Figure 1). A similar distribution of the SGI-4 sequence was observed in the whole genome sequences of 1697 *S*. Typhimurium and monophasic *S*. Typhimurium ST34 isolates from human clinical cases from England and Wales during 2014 and 2015, (Supplementary Figure 2). Of 797 *S*. Typhimurium isolates that were present outside the monophasic *S*. Typhimurium ST34 clade, just five contained the SGI-4 sequence, and these all shared a recent common ancestor with the monophasic *S*. Typhimurium ST34 clade. Six isolates from outside the monophasic *S*. Typhimurium ST34 clade contained most or all of the *sil* or *pco* genes present on SGI-4, but none also contained the SGI-4 sequence. Within the monophasic *S*. Typhimurium ST34 clade 38 isolates (4%), which were distributed sporadically throughout the clade in 28 blocks or individual leaves of the phylogenetic tree, lacked the SGI-4 sequence (Supplementary Figure 2).

### SGI-4 confers enhanced resistance to copper

A cluster of 18 ORFS on SGI-4 exhibited similarity to genes predicted to encode an RND-family efflux pump previously designated as either *cusRS cusCFBA* (17) or *silRS silCBAP* (22), and a multicopper oxidase *pcoABDRSE pcoEG (pco* locus) involved in copper and silver resistance. Alignment with sequences from homologues available in sequence databases indicated that the *cusRSCFBA* genes on the *E. coli* chromosome form an outgroup distinct from a closely-related cluster of genes that have evolved from an ancestor close to the original *silRSCBA* on pMG101 (Supplementary Figure 2). For this reason, we designated SGI-4 encoded RND-family efflux pump genes as *silRS silCBAP (sil* locus).

In order to investigate the impact of the acquisition of SGI-4 on metal ion resistance, we compared the MICs for copper sulphate of five strains of monophasic *S*. Typhimurium ST34, with three strains each of U288 and DT104 that lacked SGI-4, and a single monophasic *S*. Typhimurium ST34 strain that also lacked SGI-4 due to deletion (Figure 4). During culture in aerobic or microaerobic conditions the presence of SGI-4 had a small but significant impact on the MIC for copper. However, under anaerobic conditions, the MICs for strains lacking SGI-4 decreased by around five-fold; in contrast, SGI-4-containing monophasic *S*. Typhimurium ST34 strains had similar MICs under both aerobic and microaerobic conditions.

**Figure 4.**
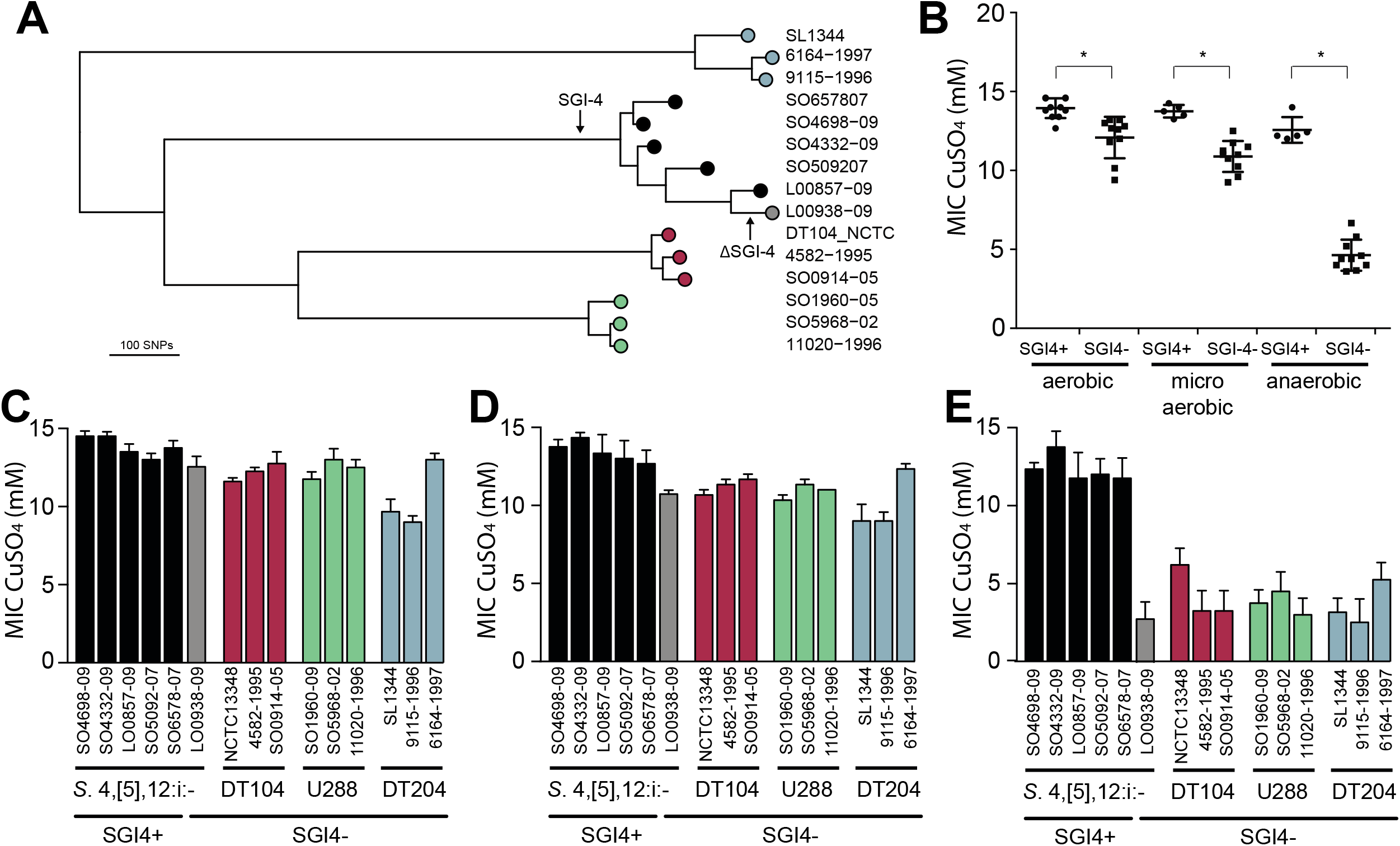
MIC for copper sulphate of representative strains of monophasic *S*. Typhimurium ST34 and *S*. Typhimurium. (A) A mid-point rooted phylogenetic tree constructed using sequence variation in the core genome with reference to *S*. Typhimurium strain SL1344 whole genome sequence (accession FQ312003). Leaves of the tree corresponding to representative strains are labelled as filled circles colour coded for SGI-4+ monophasic *S*. Typhimurium ST34 (black) and SGI-4^-^ monophasic *S*. Typhimurium ST34 (grey), and *S*. Typhimurium DT104 complex (red), U288 complex (green), and DT204 complex (blue) are shown. (B) Mean MIC for copper sulphate of SGI-4+ (filled circles) and SGI-4^-^ (filled squares) monophasic *S*. Typhimurium ST34 and *S*. Typhimurium strains during aerobic, microaerobic and anaerobic culture. (C) Bars indicate the mean MIC for copper sulphate for each strain +/− standard deviation. Colours match the tree leaf circles in (A).

### Enhanced resistance of monophasic S. Typhimurium ST34 to copper *in vitro* is mediated by the *silCBA* genes

SGI-4 encoded two clusters of genes with approximately 95% sequence identity to genes encoding an RND family efflux pump *(silCBA)*, and a multicopper oxidase system (*pcoABDE*) that have previously been implicated in resistance to copper. To determine the relative role of these two loci in copper resistance we determined the MICs of mutants of monophasic *S*. Typhimurium ST34 strain SO4698-09 that had deletions of either *silCBA*, *pcoABC* or both these loci (Figure 5). When functional *silCBA* genes were present, deletion of *pcoABDE* genes alone had no effect on resistance to copper under any of the conditions evaluated. Deletion of *silCBA* genes resulted in a small but significant (p<0.05) decrease in the MIC for copper under aerobic and microaerobic conditions, and a more substantial decrease under anaerobic conditions. However, even under anaerobic conditions, the effect of the *silCBA* deletion on the MIC for copper was not comparable to the absence of SGI-4 in U288, DT104 and DT204 isolates which had a MIC that was below 5 mg/ml copper sulphate. Copper resistance at a similar level to these strains was only observed in monophasic *S*. Typhimurium ST34 strain SO4698-09 when both *silCBA* and *pcoABDE* were deleted.

**Figure 5.**
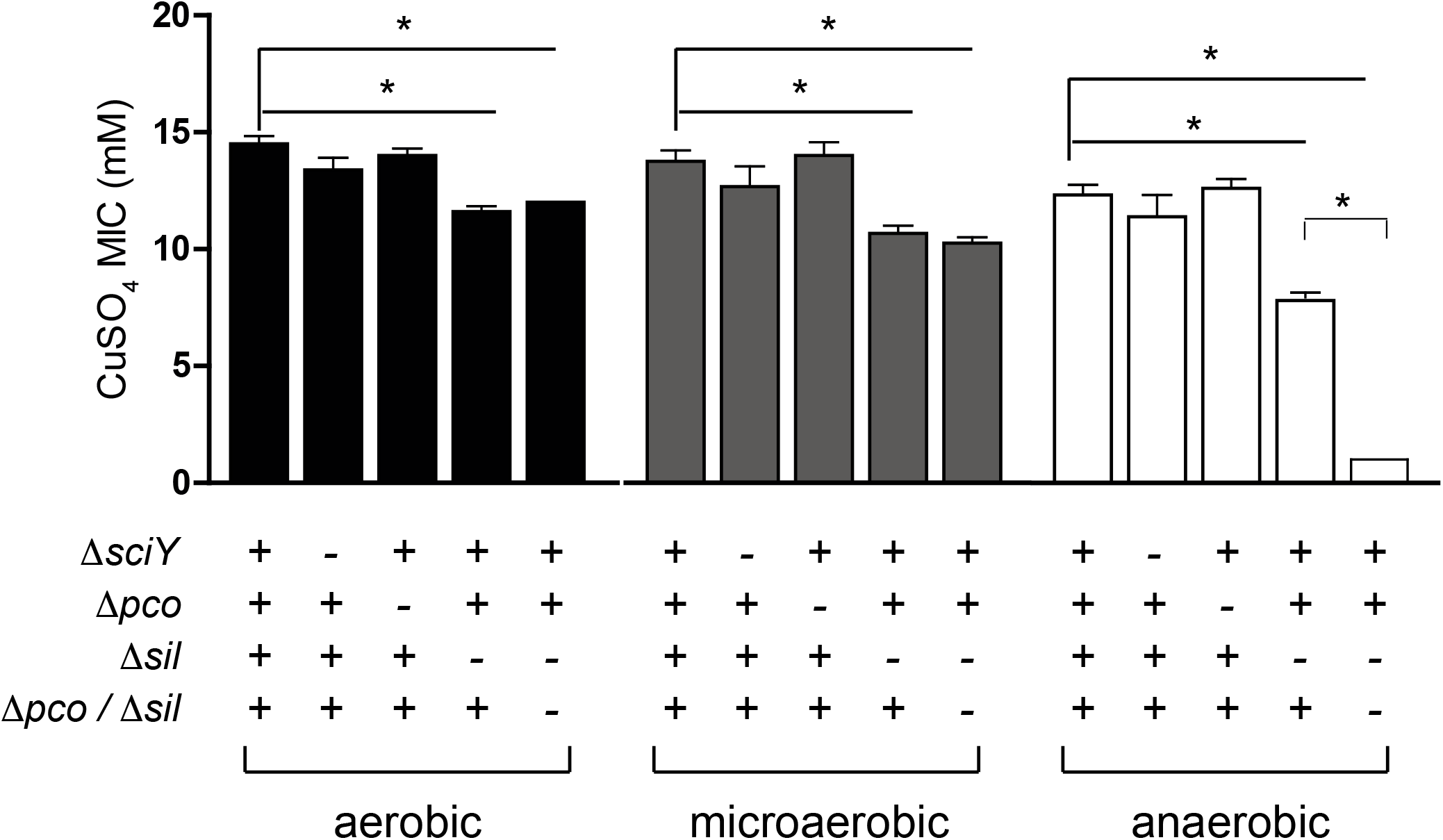
MIC for copper sulphate of monophasic S. Typhimurium ST34 strain SO4698-09 containing defined deletions in *sil* and *pco* genes. Bars indicate the mean MIC +/− standard deviation of four biological replicates during culture in either aerobic, microaerobic or anaerobic conditions. The genotype with respect to *sciY* (negative control), *pco* and *sil* are indicated as present (+) or deleted and replaced with a cat gene (-). * indicates the MIC was significantly different from wild type strain SO4698-09 (p<0.05).

### The presence of *silCBA* genes on SGI-4 alters expression of *pcoA* and the native copper homeostasis gene *copA*

To investigate any apparent redundancy of *pcoABDE* in the presence of *silCBA*, we determined the expression of *pcoA* in the presence or absence of *silCBA*. The expression of *pcoA* was not affected by increasing concentrations of copper, unless the *silCBA* genes were deleted from monophasic *S*. Typhimurium ST34 strain SO4698-09 (Figure 6). In the absence of *silCBA* induction was around ten-fold that achieved in the absence of copper in the growth medium.

**Figure 6.**
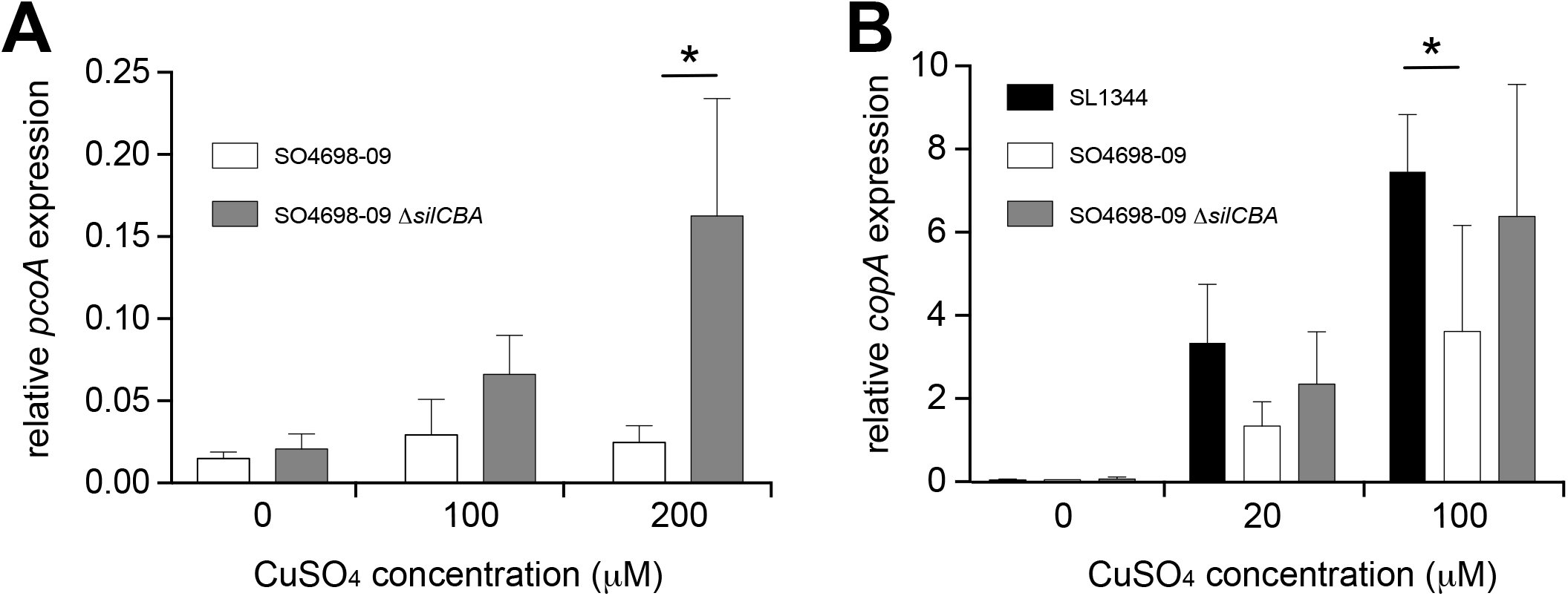
Presence of the *silCBA* genes decreases relative transcript abundance of *pcoA* and *copA*. The abundance of *pcoA* (A) or *copA* (B) transcript relative to the house keeping gene *rpoD* quantified by quantitative RT-PCR from total RNA prepared from mid-log phase cultures in LB broth supplanted with copper sulphate. Bars indicate mean relative transcript from five biological replicates +/− standard deviation.

To determine the impact of SGI-4 acquisition on typical copper homeostasis, we also investigated the expression of the *copA* gene that encodes the native ATPase protein involved in transport of copper from the cytoplasm into the periplasm in *Salmonella*. The expression of *copA* increased in a dose-dependent manner in the presence of copper in *S*. Typhimurium strain SL1344, which lacked SGI-4, and in the monophasic *S*. Typhimurium ST34 strain SO4698-09 that encoded the island. However, expression of *copA* was significantly higher in *S*. Typhimurium strain SL1344 than in monophasic *S*. Typhimurium ST34 strain SO4698-09 during culture in media with 100mM copper. Deletion of *cusACFBA* genes in strain SO4698-09 resulted in increased expression of *copA* approaching the expression level observed in SL1344 in 100 μM of CuSO4 (Figure 6).

### Presence of the *silCBA* genes does not affect SodCI-mediated resistance to macrophage killing

It is expected that expression of the *silCBA* encoded RND efflux pump would result in transport of copper from the cytoplasm and periplasm to the external milieu (29), depleting the pool of copper available in the periplasm. Our observations were consistent with this: expression of *pcoA*, a periplasmic multicopper oxidase, was not induced during culture in 100mM copper, unless the *silCBA* was deleted from SGI-4. We then tested the hypothesis that the presence of *silCBA* affected the function of SodCI, a periplasmic superoxide dismutase that uses copper as co-factor; the necessary copper is supplied by the CopA ATPase copper transporter and the periplasmic copper binding protein CueP (30). SodCI dismutates oxygen free radicals produced by phagosome associated NADPH-dependent oxidase (Phox) of macrophages infected with *Salmonella*.

To test whether altered copper homeostasis affected SodCI-mediated survival we infected gamma interferon-activated RAW macrophages with mutants of monophasic *S*. Typhimurium ST34 strain SO4698-09 that encoded either *silCBA* and *sodCI*, or lacked one or both of these loci. We then determined the change in intracellular viable counts of *S*. Typhimurium in macrophages between 2 hours and 24 hours post inoculation (Figure 7). Strain SO4698-09 exhibited a net replication of nearly ten-fold after 24 hours in RAW macrophages, and deletion of the *sodCI* gene resulted in a small but significant (p<0.05) reduction in net replication. This was consistent with previous reports that SodCI is required by the bacterium to evade the killing mechanisms of macrophages. However, deletion of the *silCBA* genes did not result in a significant decrease in net replication in RAW macrophages, but additional deletion of *sodCI* resulted in a similar decrease in net replication to that observed in the presence of the *silCBA* genes. These data are consistent with a fully functional SodCI gene in the presence of altered copper homeostasis due to the presence of the *silCBA* genes.

**Figure 7.**
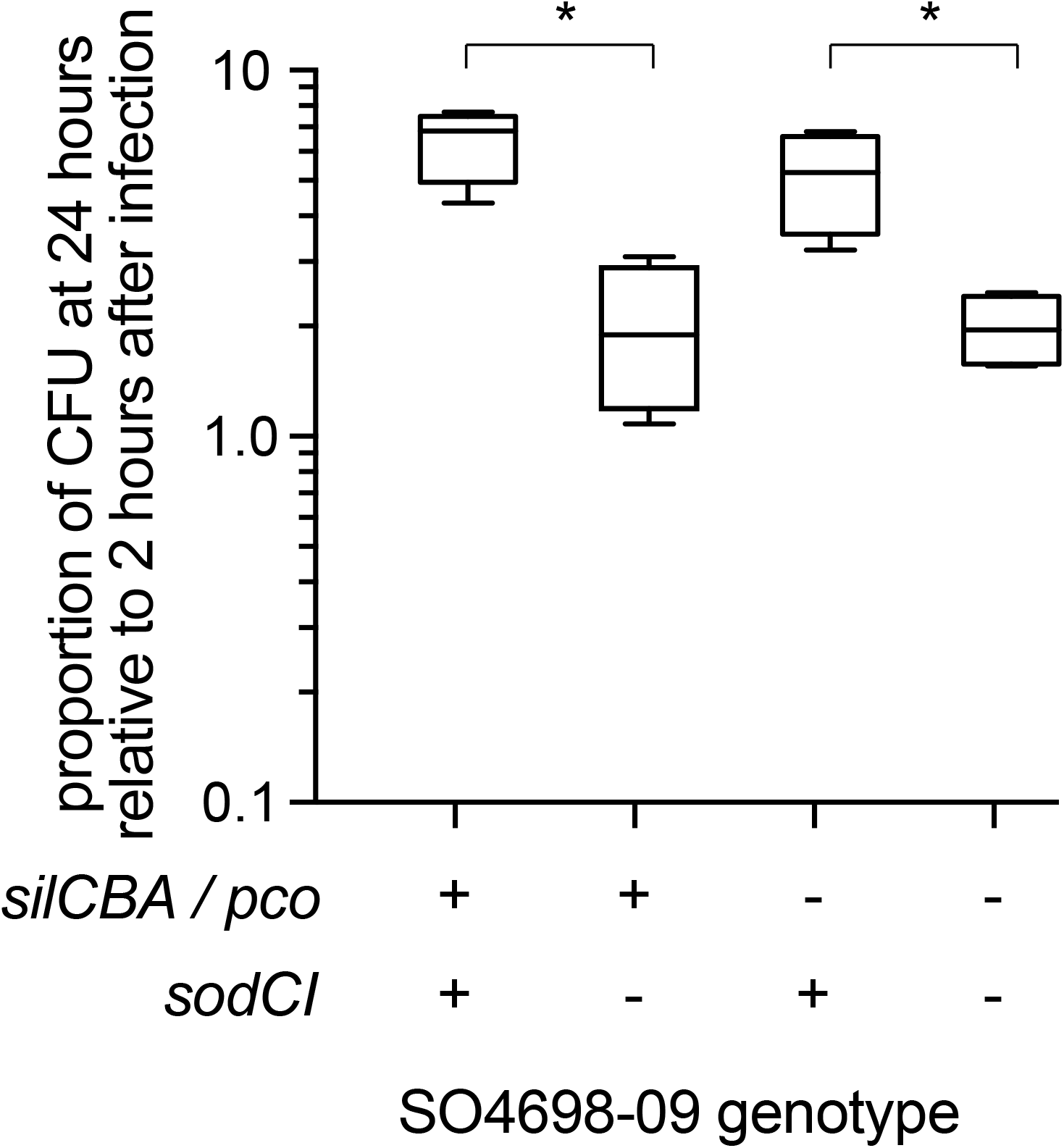
Net replication of monophasic S. Typhimurium ST34 in activated RAW macrophages is sodCI dependent but *sil* / *pco* independent. Box and whiskers plot indicating the mean number of bacterial CFUs 24 hours post infection, relative to 2 hours post infection of wild type monophasic *S*. Typhimurium strain SO4698-09 and otherwise isogenic strains containing deletions of the *sodCI* gene, the *sil/pco* genes together, or all three loci deleted.

## DISCUSSION

To achieve vertical transfer ICE undergo replication within the chromosome; for horizontal transfer they undergo excision from the genome. Horizontal transfer of ICE involves common processes including excision from the host chromsome, circularisation, coupling to a type IV secretion system (T4SS), conjugative transfer, and integration within a recipient genome at an attachment site *(attB)* (31). A number of ORFs that exhibited sequence similarity to DNA processing enzymes were identified on SGI-4. Sequences of two of these exhibited similarity to site-specific tyrosine recombinases *xerC* and *xerD*, which are predicted to mediate integration and excision. However, we were unable to identify a recombination direction factor (RDF), which are known to exhibit little sequence conservation. ORF86 had sequence similarity to *traI* that encodes a relaxase protein. This relaxase protein is capable of binding to dsDNA at the origin of transfer (oriT) and inducing a single strand nick; in conjunction with putative *uvrB* DNA helicase (ORF85) and *topB* topoisomerase (ORF 13) this facilitates unwinding of the DNA helix (32). At the left end of SGI-4, *parA* and *parB* orthologues (ORFs 1 and 5) encode putative chromosome partitioning proteins. SGI-4 also contained a number of ORFS with similarity to genes encoding components of type IV secretion systems (T4SSs) involved in conjugative transfer of DNA: ORF26 of SGI-4 encoded a putative type IV coupling protein (T4CP), TraD, which initiates conjugal transfer; ORF51 encodes a putative inner membrane protein, TraG, for T4SS stabilisation; ORF53 encodes a TraU orthologue involved in pilus assembly; ORF19 encodes a PilL F-type pilus protein; and ORF45 encoding an F-pilus assembly protein. These ORFS were present within several apparent operons (ORFs 14-27, 37-46, 48-53), consisting of multiple genes encoding proteins with no significant similarity to proteins in available databases but often associated with MGEs.

In an analysis of 1,124 taxonomically diverse complete prokaryotic genomes, 335 putative ICEs were detected based on the presence of T4SS and a relaxase gene in close proximity, indicating the widespread nature of these MGEs (31). Despite the presence of multiple genes on SGI-4 with sequence similarity to those with functions required for the transfer of ICEs, including T4SS and relaxase, SGI-4 did not exhibit extensive similarity to previously characterised mobile genetic elements. Seven complete genomes from diverse species of Enterobacteriaceae that were added to available databases since the analysis by Guglielmini et al., (31), had between 75% and 99% nucleotide sequence identity with SGI-4 and between 41% and 76% shared core sets of genes with SGI-4. As is common for ICEs, core genes were interspersed with cargo genes with functions that were unrelated to mobilisation but had the potential to modify the phenotype of the host bacterium (27). In particular, genes involved in resistance to heavy metals were present on SLEs, with five containing at least one locus associated with resistance to copper, silver, mercury or arsenic Two SLEs from *Erwinia tracheiphila* and *Edwardsiella ictaluri* were particularly closely related to SGI-4, but contained unrelated cargo genes, highlighting the rapid divergence achieved by horizontal gene transfer in this family of ICE. Our analysis is consistent with SGI-4 evolving from a common ancestor of these ICEs by acquisition of *ars, sil* and *pco* cargo genes involved in resistance to metal ions.

With the exception of a small outbreak in a burns unit associated with a *S*. Typhimurium strain in the 1970s, historically, the *sil* and *pco* loci encoding resistance to copper and silver, have rarely been associated with *Salmonella enterica* isolates. However, *sil* and *pco* genes, have been associated with three lineages of monophasic *S*. Typhimurium in the past 20 years: the ‘European clone’ (strain ST34); the ‘Spanish clone’; and the ‘Southern European clone’ (11). However, only 74 and 26% of the latter two, respectively, encode the *sil* and/or the *pco* genes and these were plasmid-borne, suggesting that they may be lost relatively frequently. In contrast, monophasic *S*. Typhimurium ST34 encodes *silCBA* and *pco* genes on the chromosome, and although on a mobile genetic element, we found these genes in 96% of the 797 clinical isolates from England and Wales that we evaluated. Furthermore, loss of SGI-4 and the *silCBA* and *pco* genes was sporadic, and mostly as singleton taxa on the phylogenetic tree, consistent with the absence of selection for their loss.

Given the general paucity of genes encoding copper/silver RND efflux pumps in *Salmonella enterica* and the evolutionary history of the genus, the apparent lack of a selective advantage for the loss of *silCBA* genes may indicate an important role for these genes in the monophasic *S*. Typhimurium ST34 clone. Non-functional remnants of the *cusCFBA* genes are present in a region of the genome of *S*. Typhimurium that has synteny with the genomic region of the *cusCFBA* genes of *E. coli*, indicating that they were likely to have been lost through deletion since these species shared a common ancestor (19, 33). Deletion of *cusCFBA* in the *Salmonella* ancestor is likely to have had a profound effect on copper distribution in the cell because *Salmonella* has no alternative mechanism to remove copper from cells. While *E. coli* can remove copper from the cell entirely, *Salmonella* transports copper from the cytoplasm, where it is especially toxic, to the periplasm, where it is detoxified by the copper oxidase CueO or bound to the copper binding protein, CueP, the major reservoir of copper in *Salmonella* (34). The evolution of *Salmonella* pathogenesis must, therefore, have proceeded in the context of fundamentally altered copper homeostasis. The reintroduction of the *silCBA* locus on SGI-4 did indeed alter the copper homeostasis of *Salmonella* as indicated by the lack of *pcoA* expression in the presence of the RND-family of efflux genes, which are regulated in response to copper levels in the periplasm. The consequences of altered copper homeostasis in monophasic *S*. Typhimurium ST34 is not clear but, at least in RAW macrophages *in vitro*, the dependency of SodCI copper as a cofactor did not affect resistance to macrophage killing.

Strong selection pressure may be important for retention of SGI-4 since its presence profoundly alters copper homeostasis. Monophasic *S*. Typhimurium ST34 is primarily associated with pigs, and it has been suggested that the success of this clone may, in part, have been driven by the extensive use of copper as a growth promotor in pig rearing (11, 25). Consistent with this idea, the presence of SGI-4 is correlated with enhanced resistance to copper; this appears to be particularly apparent under anaerobic conditions which are similar to those encountered by *Salmonella* in the host intestinal tract. *Salmonella* strains that lacked SGI-4, exhibited MICs for copper of around 2-3mM under anaerobic conditions, which is approximately 15% of the values achieved under aerobic conditions. Importantly, this is in the range of the concentrations of copper found in pig manure from animals that had been fed on a copper-supplemented diet (35). The presence of SGI-4, however, increased the MIC for copper (under anaerobic conditions) by approximately 500%, elevating it to levels above the concentrations likely to be encountered on pig farms. The *sil* genes encoded on SGI-4 were entirely responsible for the observed increase in MICs for copper, since deletion of *pco* genes alone did not result in a decrease in the MICs. The *pco* locus encodes a multicopper oxidase which is thought to detoxify the more damaging Cu+ to Cu2+ ions by oxidation (36). Monophasic *S*. Typhimurium ST34 encodes a native copper oxidase, CueO, and the presence of this protein may, in part, mask the activity of the *pco* locus. However, the *pco* genes were capable of enhancing resistance to copper in the absence of *cusCFBA*. Furthermore, since there was no evidence for loss of *pco* genes during clonal expansion of the monophasic *S*. Typhimurium ST34 clade, it remains possible that the Pco system may play a more prominent role in copper resistance under environmental conditions that we did not evaluate in our *in vitro* experiments.

## MATERIAL AND METHODS

### Bacterial strains and growth conditions

*S*. Typhimurium strain SL1344 was isolated from a calf in 1973 as described previously (33). Monophasic *S*. Typhimurium ST34 strains (SO4698-09, SO4332-09, LO0857-09, SO5092-07, SO6578-07, LO938-09); *S*. Typhimurium U288 strains SO1960-05, SO5968-02 and 11020-1996; *S*. Typhimurium DT104 strains (NCTC13348, 4582-1995 and SO0914-05); and two strains that were closely related to strain SL1344 (9115-1996 and 6164-1997) were isolated from various host species and their whole genome sequence determined as described previously (25, 37). Strains of *S*. Typhimurium or monophasic *S*. Typhimurium in which specific genes were replaced with either the *cat* gene conferring chloramphenicol resistance or the *aph* gene conferring resistance to kanamycin, were constructed using allelic exchange methodology as described previously (38). Briefly, pairs of oligonucleotide primers specific for amplification of the *cat* gene or the *aph* gene from plasmids pKD3 or pKD4 were synthesised with 50 nucleotide sequences on the 5’ end that were identical to the sequences immediately proximal to the ATG start and distal to the stop codon of each gene targeted for deletion. Bacterial strains were routinely cultured in Luria Bertani broth (Oxoid) supplemented with 0.03 mg/l chloramphenicol or 0.05 mg/l kanamycin as appropriate.

### Sequence analysis

A gene model for SGI-4 was constructed using Prokka (39) with a minimum ORF size of 100. Annotation of SGI-4 was achieved by aligning open reading frame (ORF) sequences to sequences in the NCBI database using BLASTn to identify genes with the greatest sequence identity. Phylogenetic trees from whole genome sequence data were constructed and single nucleotide polymorphisms (SNPs) identified in the whole genome sequences by aligning reads using BWA MEM (40), variant calling with Freebayes (41) and SNP filtering using vcflib/vcftools (42), using Snippy v3.0 (https://github.com/tseemann/snippy). Maximum-likelihood trees were constructed using a general time-reversible (GTR) substitution model with gamma correction for amongst-site rate variation with RAxML v8.0.20 and 1000 bootstraps (43). Representative isolates of *S*. Typhimurium phage types have been described previously (25), and whole genome sequences of *S*. Typhimurium isolates from routine clinical surveillance by Public Health England are available in public databases (Supplementary Table I). For reconstruction of the phylogeny of SLEs, putative SLEs were identified in genome assemblies in the NCBI non-redundant sequence database and aligned to SGI-4 excluding the *ars*, *sil* and *pco* loci, using BLASTn. Nucleotide sequences from *Edwardsiella ictaluri* (accession CP001600, 326500..380800), *Erwinia tracheiphila* (accession CP013970, 1856073..1916737), *Enterobacter cloacae* (accession CP012162, 4040884..4152906), *Enterobacter hormaechei* (accession CP010376, 3932000..4028800), *Enterobacter hormaechei* (accession CP012165, 395200..536000), *Pluralibacter gergoviae* (accession CP009450, 1604785..1778603) and *Salmonella* Cubana (accession CP006055, 4214040..4311200), were aligned using clustalW to determine the extent of similarity and sequence identity in conserved regions. A maximum likelihood tree was constructed with nucleotide sequence alignment using RAxML as described previously (43). The relationship of SLE’s was also investigated by determining the proportion of each SLE that aligned by carrying out a pair-wise comparison with discontinuous megaBLAST (44). Each SLE was used as the query sequence against each SLE as the subject to determine the percent of sequence tha aligned and the mean percent nucleotide sequence identity of the aligned sequence.

### Determination of transfer of SGI-4 *in vitro*

In order to provide a convenient selection for the presence of SGI-4, we constructed a strain of monophasic *S*. Typhimurium SO4698-09 in which the *arsC* gene on SGI-4 was replaced by the *cat* gene (SO4698-09 SGI-4 Δ*arsC::cat*), conferring resistance to chloramphenicol. To provide a selection for the recipient strain, we constructed a strain of *S*. Typhimurium SL1344 in which the36 *copA* gene was deleted and replaced by an *aph* gene, conferring resistance to kanamycin. Donors and recipients were cultured in LB broth for 18 hours at 37°C with shaking. The OD600nm of each culture was adjusted to 0.1 with fresh LB broth and 2.5 ml of each added to a 50ml tube and incubated for 18 hours at 37°C with shaking. The number of CFUs per ml of donors and recipients were quantified by culturing serial dilutions on LB agar supplemented with 0.03 mg/ml chloramphenicol or 0.05 mg/ml kanamycin, respectively. The presence of SGI-4 Δ*arsC::cat* in recipient strains was quantified by serial dilution on LB agar supplemented with chloramphenicol and kanamycin. Transfer frequency was defined as the number of recipients containing SGI-4 Δ*arsC::cat* as a proportion of donor cells in the culture. To determine whether trans-conjugant recipient strains contained SGI-4 in the same chromosomal location as the donor, the predicted right junction was amplified using specific primers spanning the left 5’-gtccctcaagtaagggaac-3’ and right 5’-aatgatcggatcttctgatgga-3’ ends of SGI-4 by PCR. The sequence of the amplicon was determined by Sanger sequencing using the same primers (Eurofin sequencing service).

### Determination of MIC for copper sulphate

Bacterial strains were cultured for 18 hours in LB broth at 37°C with shaking. To this 0.2 ml aliquots of LB broth buffered with 25mM HEPES pH7 containing a range of concentrations of copper sulphate were added to each well of a polystyrene 96-well plate and equilibrated for 24 hours at 37°C normal atmospheric, microaerobic (10% CO2, 5% H_2_, 5% O_2_ and 80% N_2_) or anaerobic (10% CO_2_, 10% H_2_ and 80% N_2_) conditions. Each well of polystyrene 96-well plates (Nunc) was inoculated with 1x10^7^ CFUs of bacterial culture containing a range of metal ions. 96-well plates were incubated at 37°C for 24 hours, and the OD600 of each well measured using a BIORAD Benchmark Plus microplate spectrophotometer. The MIC was defined as the mean concentration of copper sulphate for which the OD600 of the culture was <0.2 of from four biological replicates.

### Quantitative real time PCR expression analysis

Expression relative to transcript abundance a constitutively expressed housekeeping gene was determined as previously described (45). Total RNA was extracted from 2 ml samples from overnight cultures grown in LB to mid-log phase (OD600nm of 0.2) with aeration or in an anaerobic atmosphere (10% CO_2_, 10% H_2_, N_2_) and either with or without CuSO4 supplementation. Bacteria were harvested by centrifugation and RNA prepared using a FastRNA^TM^ spin kit for microbes (MPBio) according to the manufacturer’s instructions. RNA was treated with a TURBO DNA-free^TM^ kit (ambion, life technologies™) according to the manufacturer’s instructions before being reverse-transcribed using the QuantitTect^®^ Reverse Transcription kit (Qiagen^®^). The resulting cDNA or serial dilutions of a known quantity of genomic DNA for generation of standard curves were amplified using a QuantiFast^®^ SYBR^®^ Green PCR kit (Qiagen^®^) with specific primers 5’-GCGGACGCGTTAATTGAAAC-3’ and 5’-TGTTGGCTTTCTTCATCGGC-3’ for *copA*; 5’-GAATGGACCACAGCCAGATG-3’ and 5’-GACGCAGGATGACTTTGCAT-3’ for *pcoA*; and 5’-GTCAACAGTATGCGCGTGAT-3’ and 5’-GATAGCGGCATTGAACCAGG-3’ for *rpoD* housekeeping using the Applied Biosystems^®^ 7500 real-time PCR system. The expression of *copA* or *pcoA* is presented relative to transcript abundance of the *rpoD* gene.

### RAW264.7 macrophage survival assay

Murine macrophages (RAW 264.7, ATCC, Rockville, MD) were grown in minimum essential medium (Sigma Aldrich) supplemented with 10 % foetal bovine serum, L-glutamine (2 mM) and 1 x nonessential amino acids. For infection studies, 2x10^5^ murine cells per well were seeded into 24-well plates 48 h before infection. After 24 h the murine cells were activated with 50 ng/ml of IFN-γ in the presence or absence of 125 μM CuSO_4_ for a further 24 h. Cells were infected with bacteria in stationary phase culture at a multiplicity of infection of 20. Plates were centrifuged at 1500 rpm for 3 minutes followed by incubation of the cells for 30 minutes. The medium was then exchanged for fresh medium of the same composition with the exception of the addition of gentamycin (100 μg/ml). After either 2 or 24 h, the cells were washed with PBS, lysed with PBS 1% Triton and the number of bacterial CFUs determined by culturing serial dilutions on LB agar. The data were expressed as the proportion of cfu determined at 24 h relative to the cfu determined at 2 h.

### Statistical analysis

The Mann-Whitney U test was used to test the null hypothesis that randomly selected values in a sample were equally likely to be greater or smaller than from a second sample, using a false discovery rate of 0.05 to reject the null hypothesis.

## ACKNOWLEDGEMENTS

RK was supported by research grants BB/R012504/1, BB/N007964/1 and BB/M025489/1 from the BBSRC. OC was supported by a BBSRC DTP studentship (BB/M011216/1).

---

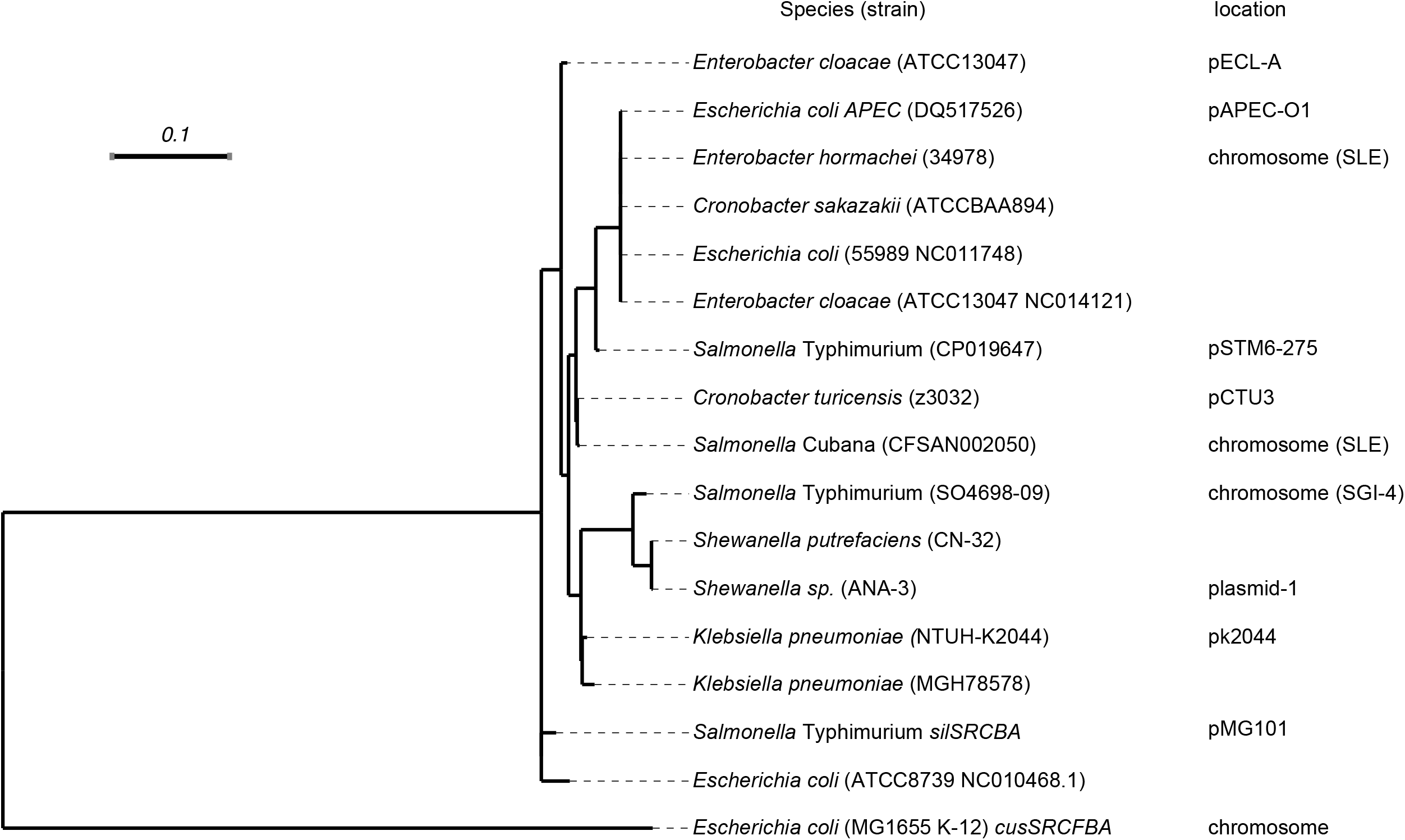

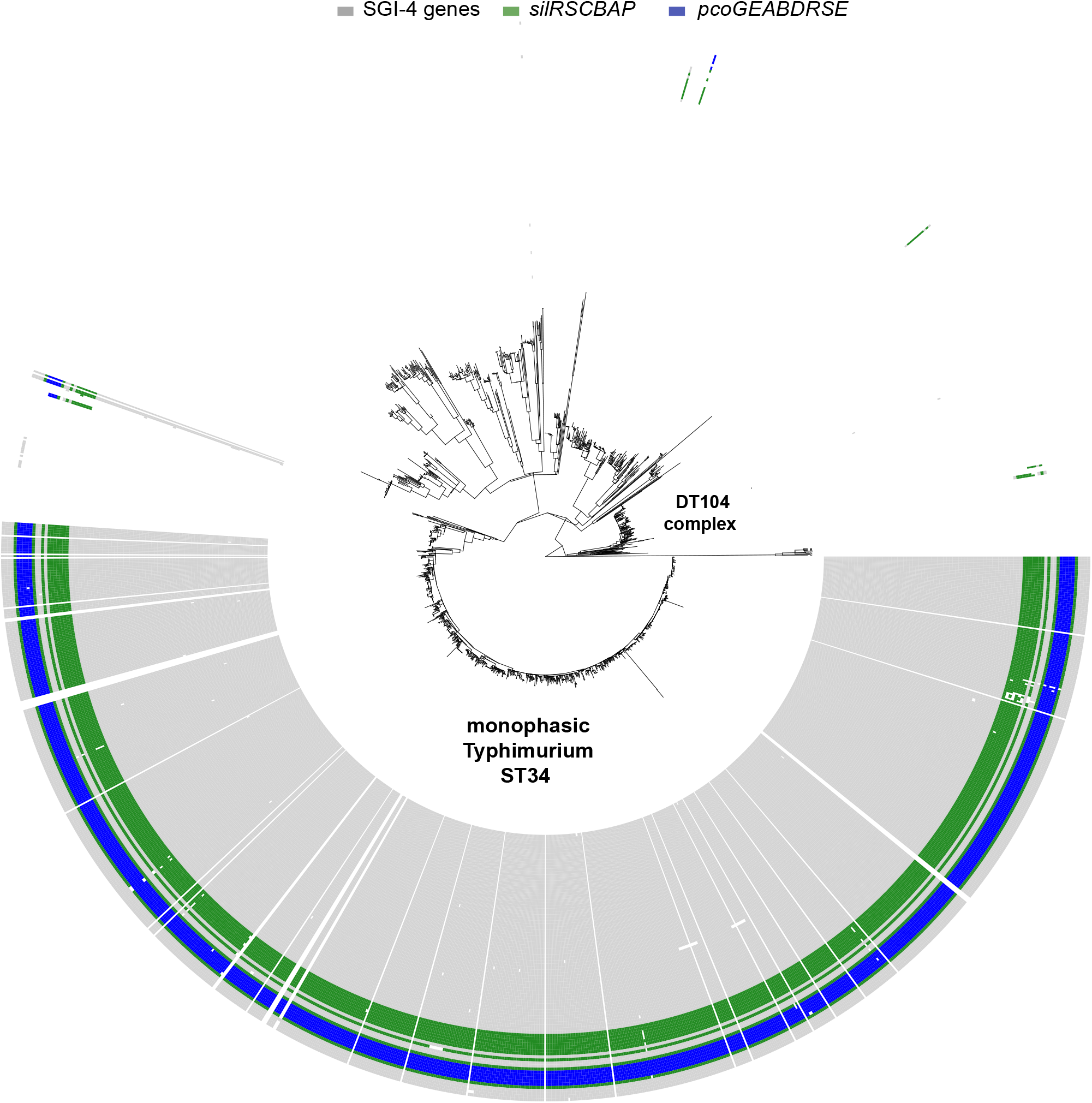

## REFERENCES

1. Hugas M, Beloeil P. Controlling Salmonella along the food chain in the European Union-progress over the last ten years. Euro surveillance : bulletin Europeen sur les maladies transmissibles = European communicable disease bulletin. 2014;19(19).

2. Branchu P, Bawn M, Kingsley RA. Genome variation and molecular epidemiology of Salmonella Typhimurium pathovariants. Infect Immun. 2018;86(8):e00079–18.

3. Rabsch W. Salmonella Typhimurium Phage Typing for Pathogens. In: Schatten H, Eisenstark A, editors. Salmonella, Methods and Protocols. Methods in Molecular Biology. 394 ed.Totowa, New Jersey: Humana Press; 2007. p. 177–212.

4. Rabsch W, Tschape H, Baumler AJ. Non-typhoidal salmonellosis: emerging problems. Microbes Infect. 2001;3(3):237–47.

5. Threlfall EJ. Epidemic Salmonella typhimurium DT 104-a truly international multiresistant clone. Journal of Antimicrobial Chemotherapy. 2000;46(1):7–10.

6. Andres-Barranco S, Vico JP, Marin CM, Herrera-Leon S, Mainar-Jaime RC. Characterization of Salmonella enterica Serovar Typhimurium Isolates from Pigs and Pig Environment-Related Sources and Evidence of New Circulating Monophasic Strains in Spain. J Food Prot. 2016;79(3):407–12.

7. Arguello H, Sorensen G, Carvajal A, Baggesen DL, Rubio P, Pedersen K. Prevalence, serotypes and resistance patterns of Salmonella in Danish pig production. Res Vet Sci. 2013;95(2):334–42.

8. Hauser E, Tietze E, Helmuth R, Junker E, Blank K, Prager R, et al. Pork contaminated with Salmonella enterica serovar 4,[5],12:i:-, an emerging health risk for humans. Appl Environ Microbiol. 2010;76(14):4601–10.

9. Bonardi S. Salmonella in the pork production chain and its impact on human health in the European Union. Epidemiol Infect. 2017;145(8):1513–26.

10. Hopkins KL, de Pinna E, Wain J. Prevalence of Salmonella enterica serovar 4, [5],12:i:-in England and Wales, 2010. Eurosurveillance. 2012;17(37):10–6.

11. Mourao J, Novais C, Machado J, Peixe L, Antunes P. Metal tolerance in emerging clinically relevant multidrug-resistant Salmonella enterica serotype 4,[5],12:i:-clones circulating in Europe. Int J Antimicrob Agents. 2015;45(6):610–6.

12. Antunes P, Mourao J, Pestana N, Peixe L. Leakage of emerging clinically relevant multi drug-resistant Salmonella clones from pig farms. J Antimicrob Chemother. 2011;66(9):2028–32.

13. Macomber L, Imlay JA. The iron-sulfur clusters of dehydratases are primary intracellular targets of copper toxicity. Proc Natl Acad Sci U S A. 2009;106(20):8344–9.

14. Rensing C, Grass G. Escherichia coli mechanisms of copper homeostasis in a changing environment. FEMS Microbiol Rev. 2003;27(2-3):197–213.

15. Franke S, Grass G, Rensing C, Nies DH. Molecular analysis of the copper-transporting efflux system CusCFBA of Escherichia coli. J Bacteriol. 2003;185(13):3804–12.

16. Grass G, Rensing C. CueO is a multi-copper oxidase that confers copper tolerance in Escherichia coli. Biochem Biophys Res Commun. 2001;286(5):902–8.

17. Outten FW, Huffman DL, Hale JA, O’Halloran TV. The independent cue and cus systems confer copper tolerance during aerobic and anaerobic growth in Escherichia coli. J Biol Chem. 2001;276(33):30670–7.

18. Brown NL, Barrett SR, Camakaris J, Lee BT, Rouch DA. Molecular genetics and transport analysis of the copper-resistance determinant (pco) from Escherichia coli plasmid pRJ1004. Mol Microbiol. 1995;17(6):1153–66.

19. McClelland M, Sanderson KE, Spieth J, Clifton SW, Latreille P, Courtney L, et al. Complete genome sequence of Salmonella enterica serovar Typhimurium LT2. Nature. 2001;413(6858):852–6.

20. Pontel LB, Soncini FC. Alternative periplasmic copper-resistance mechanisms in Gram negative bacteria. Mol Microbiol. 2009;73(2):212–25.

21. Espariz M, Checa SK, Audero ME, Pontel LB, Soncini FC. Dissecting the Salmonella response to copper. Microbiology. 2007;153(Pt 9):2989–97.

22. Gupta A, Matsui K, Lo J-F, Silver S. Molecular basis for resistance to silver cations in Salmonella. Nature medicine. 1999;5(2):183.

23. McHugh GL, Moellering RC, Hopkins CC, Swartz MN. Salmonella typhimurium resistant to silver nitrate, chloramphenicol, and ampicillin. Lancet. 1975;1(7901):235–40.

24. Campos J, Mourao J, Marcal S, Machado J, Novais C, Peixe L, et al. Clinical Salmonella Typhimurium ST34 with metal tolerance genes and an IncHI2 plasmid carrying oqxAB-aac(6’)-Ib-cr from Europe. J Antimicrob Chemother. 2016;71(3):843–5.

25. Petrovska L, Mather AE, AbuOun M, Branchu P, Harris SR, Connor T, et al. Microevolution of Monophasic Salmonella Typhimurium during Epidemic, United Kingdom, 2005-2010. Emerg Infect Dis. 2016;22(4):617–24.

26. Hochhut B, Waldor MK. Site-specific integration of the conjugal Vibrio cholerae SXT element into prfC. Mol Microbiol. 1999;32(1):99–110.

27. Wozniak RA, Waldor MK. Integrative and conjugative elements: mosaic mobile genetic elements enabling dynamic lateral gene flow. Nature Reviews Microbiology. 2010;8(8):552.

28. Li X, Xie Y, Liu M, Tai C, Sun J, Deng Z, et al. oriTfinder: a web-based tool for the identification of origin of transfers in DNA sequences of bacterial mobile genetic elements. Nucleic acids research. 2018.

29. Gupta A. Multidrug-resistant typhoid fever in children: epidemiology and therapeutic approach. Pediatr Infect Dis J. 1994;13(2):134–40.

30. Osman D, Patterson CJ, Bailey K, Fisher K, Robinson NJ, Rigby SE, et al. The copper supply pathway to a Salmonella Cu,Zn-superoxide dismutase (SodCII) involves P(1B)-type ATPase copper efflux and periplasmic CueP. Mol Microbiol. 2013;87(3):466–77.

31. Guglielmini J, Quintais L, Garcillán-Barcia MP, De La Cruz F, Rocha EP. The repertoire of ICE in prokaryotes underscores the unity, diversity, and ubiquity of conjugation. PLoS genetics. 2011;7(8):e1002222.

32. Salyers AA, Shoemaker NB, Stevens AM, Li L-Y. Conjugative transposons: an unusual and diverse set of integrated gene transfer elements. Microbiological reviews. 1995;59(4):579–90.

33. Kroger C, Dillon SC, Cameron AD, Papenfort K, Sivasankaran SK, Hokamp K, et al. The transcriptional landscape and small RNAs of Salmonella enterica serovar Typhimurium. Proc Natl Acad Sci U S A. 2012;109(20):E1277–86.

34. Osman D, Waldron KJ, Denton H, Taylor CM, Grant AJ, Mastroeni P, et al. Copper homeostasis in Salmonella is atypical and copper-CueP is a major periplasmic metal complex. J Biol Chem. 2010;285(33):25259–68.

35. Nicholson FA, Chambers BJ, Williams JR, Unwin RJ. Heavy metal contents of livestock feeds and animal manures in England and Wales. Bioresource Technol. 1999;70(1):23–31.

36. Lee SM, Grass G, Rensing C, Barrett SR, Yates CJ, Stoyanov JV, et al. The Pco proteins are involved in periplasmic copper handling in Escherichia coli. Biochem Biophys Res Commun. 2002;295(3):616–20.

37. Mather AE, Lawson B, de Pinna E, Wigley P, Parkhill J, Thomson NR, et al. Genomic Analysis of Salmonella enterica Serovar Typhimurium from Wild Passerines in England and Wales. Appl Environ Microbiol. 2016;82(22):6728–35.

38. Datsenko KA, Wanner BL. One-step inactivation of chromosomal genes in Escherichia coli K-12 using PCR products. Proc Natl Acad Sci U S A. 2000;97(12):6640–5.

39. Seemann T. Prokka: rapid prokaryotic genome annotation. Bioinformatics. 2014;30(14):2068–9.

40. Li H. Aligning sequence reads, clone sequences and assembly contigs with BWA-MEM. 2013.

41. Garrison E, Marth G. Haplotype-based variant detection from short-read sequencing 2012[

42. Danecek P, Auton A, Abecasis G, Albers CA, Banks E, DePristo MA, et al. The variant call format and VCFtools. Bioinformatics. 2011;27(15):2156–8.

43. Stamatakis A. RAxML-VI-HPC: maximum likelihood-based phylogenetic analyses with thousands of taxa and mixed models. Bioinformatics. 2006;22(21):2688–90.

44. Morgulis A, Coulouris G, Raytselis Y, Madden TL, Agarwala R, Schaffer AA. Database indexing for production MegaBLAST searches. Bioinformatics. 2008;24(16):1757–64.

45. Branchu P, Matrat S, Vareille M, Garrivier A, Durand A, Crepin S, et al. NsrR, GadE, and GadX interplay in repressing expression of the Escherichia coli O157:H7 LEE pathogenicity island in response to nitric oxide. PLoS Pathog. 2014;10(1):e1003874.

